# Genomic Data from an Endangered Amphibian Reveal Unforeseen Consequences of Fragmentation by Roads

**DOI:** 10.1101/306340

**Authors:** Evan McCartney-Melstad, Jannet K. Vu, H. Bradley Shaffer

## Abstract

Roads fragment landscapes and can cause the loss of metapopulation dynamics in threatened species, but as relatively new landscape features, few studies have had the statistical power to genetically examine road effects. We used DNA sequence data from thousands of nuclear loci to characterize the population structure of New York-endangered Eastern tiger salamanders *(Ambystoma tigrinum)* on Long Island and quantify the impacts of roads on population fragmentation. We uncovered highly genetically structured populations over an extremely small spatial scale (approximately 40 km^2^) in an increasingly human-modified landscape. Geographic distance and the presence of roads between ponds are both strong predictors of genetic divergence, suggesting that both natural and anthropogenic factors are responsible for the observed patterns of genetic variation. Our study demonstrates the value of genomic approaches in molecular ecology, as these patterns did not emerge in an earlier study of the same system using microsatellite loci. Ponds supported small effective population sizes, and pond surface area showed a strong positive correlation with salamander population size. When combined with the high degree of structuring in this heavily modified landscape, our study indicates that these endangered amphibians require management at the individual pond, or pond cluster, level. Particular efforts should be made to preserve large vernal pools, which harbor the greatest genetic diversity, and their surrounding upland habitat. Contiguous upland landscapes between ponds that facilitate natural metapopulation connectivity and demographic rescue from future local extirpations should also be protected.

## Introduction

Biological conservation often occurs at small spatial scales and in human dominated landscapes including the urban-wildland interface, agricultural habitat mosaics, and small urban parklands. Critical management questions frequently come down to the valuation of remnant habitat patches, the impact of roads or other anthropogenic disturbances on those patches, and the viability of small, fragmented populations. Genetic, and, increasingly, genomic analyses constitute a powerful tool kit for conservation evaluation and action, particularly for secretive species such as reptiles and amphibians (Shaffer et al., 2015). When studying endangered species, conservationists often focus on the value of small protected populations or the impact of a road between nearby populations that historically exchanged migrants, tracking these changes over short temporal scales. As conservation and population biologists, we are generally most interested in dynamics across a few kilometers on specific landscapes, rather than the larger spatiotemporal scales spanning thousands of kilometers and generations that are typically the purview of population genetics.

Particularly for secretive, cryptic, low-vagility species, including amphibians, reptiles, small mammals, and many invertebrates that often move a kilometer or less per generation (Blaustein, Wake, & Sousa, 1994), acquiring the data needed for effective management and trend analysis is virtually impossible with traditional ecological methods. Indirect inferences derived from population genetic and genomic datasets constitute the most effective way forward for these taxa, and those approaches have continued to expand in both influence and sophistication in recent decades. Establishing the number of salamanders utilizing a breeding site or assessing the fragmentation effects of a road on burying beetles may require decades of intensive field work, but estimating the effective population size of that pond or genetic differentiation across that road can potentially be accomplished with a single field trip and a few months of lab work. For protected or endangered species, even rough estimates of key population parameters allow resource managers to parameterize population viability analyses (PVAs) and make scientifically sound, rapid assessments that are key for effective management.

A critical consideration when using molecular ecological methods to detect trends and parameterize models at very fine spatial scales has always been their limits of resolution. Theory and decades of empirical study have established that populations in close proximity tend to be very closely related (Wright, 1943), genetic change accumulates over time, and the ability to detect differentiation between genetically very closely related populations is limited by the number of samples and genetic loci assayed (Felsenstein, 2006; Patterson, Price, & Reich, 2006). Until recently, most population genetic studies of high conservation value species, and virtually all studies of amphibians have been limited to mitochondrial DNA or a small number of nuclear loci (typically microsatellites), although this is slowly changing as genomic technologies have evolved and been applied to amphibians (Keinath, Voss, Tsonis, & Smith, 2016; McCartney-Melstad, Gidiş, & Shaffer, 2017; McCartney-Melstad, Mount, & Shaffer, 2016; C. E. Newman & Austin, 2016; Portik, Smith, & Bi, 2016). However, most systems that might benefit from genomic scale data have yet to receive the attention of conservation genomicists.

Here, we present a comparative case study of the Eastern tiger salamander (*Ambystoma tigrinum*) to address two key questions that have broad implications across taxa. First, over the small spatial scales that characterize many conservation problems, does the added resolution of genomic-scale data make a substantive difference in the inferences and parameter estimates necessary for effective management? Second, when a well-designed microsatellite study is repeated with independent genome-level data from the same landscape, do the resulting conservation outcomes change?

The Eastern tiger salamander is a New York-listed endangered species (6 CRR-NY 182.5) that was historically found in few scattered localities across eastern New York including Albany and Rockland Counties, and across Long Island in sandy, vernal pool habitats. The species has experienced dramatic declines in the region, and it is currently restricted to Suffolk County (Bishop, 1941; Stewart & Rossi, 1981; Breisch, pers. comm.), with approximately 90 remaining breeding ponds in central Long Island (New York State Department of Environmental Conservation, 2015). The species suffers a range of threats including disease, pollution, predation by invasive species, climate change-induced sea level rise, habitat loss, road mortality, illegal collecting and population fragmentation (Titus, Bell, Becker, & Zamudio, 2014). Telemetry studies documented individuals traveling at least 500 meters from breeding ponds and confirmed that they tend to avoid paved roads, dirt roads, and grassy areas (Madison & Farrand, 1998).

Prior genetic work using twelve microsatellite loci recovered two population clusters of *A. tigrinum* across 17 ponds spanning 50 km on Long Island, both of which exhibited low diversity and high relatedness among ponds (Titus et al., 2014). The authors attributed the low diversity and high relatedness to a combination of post-glacial colonization from North Carolina (Church, Kraus, Mitchell, Church, & Taylor, 2003) and relatively frequent migration of salamanders between breeding ponds. Their primary conclusion was that endangered Long Island tiger salamanders were genetically uniform but differentiated from the nearest populations in New Jersey approximately 250 km distant. Most of the 17 Long Island ponds analyzed by Titus *et al.* (2014) were fewer than six kilometers apart, providing a challengingly small landscape over which to make inferences. The microsatellite loci showed relatively low diversity (1–13 alleles per locus across ponds and an average of 1–3 alleles per locus within ponds), and therefore were not as variable as is typical for these markers (Reyes-Valdés, 2013).

The Titus *et al.* (2014) research is an example of much of the best work on amphibian conservation genetics: they sampled a representative set of breeding ponds each for a large population sample, used a reasonable number of variable genetic markers, and drew conclusions based on those data about population connectivity, fragmentation, and effective breeding size. However, their study also confronted empirical roadblocks. The microsatellites exhibited low levels of variation, resulting in reduced statistical power and the somewhat surprising result that gene flow among these low-vagility, pond-breeding animals was consistently high. Whether that inference represents the actual biology of the species, or is an artifact of the genetic tools available to their study, is an open question that speaks to the broader interpretation of similar studies (e.g. Jehle, Burke, & Arntzen, 2005; Lampert, Rand, Mueller, & Ryan, 2003; R. A. Newman & Squire, 2001; Zamudio & Wieczorek, 2007).

To explore this question, we applied a genomic target capture approach with 5,237 random nuclear exons to tiger salamanders sampled from the same set of ponds in central Long Island. We sought to answer four questions: 1) To what degree are ponds genetically connected to or differentiated from one another?, 2) What are the effective population sizes of ponds in the system, are they related to pond area, and how do these values compare to other amphibians?, 3) What are the effects of roads and traffic patterns on connectivity between ponds in the system?, and 4) Does the increased resolution of the genomic dataset provide additional insights for conservation above beyond those derived from microsatellites? Our results indicate that the gain in using genomic data is substantial, arguing that it may well be worth the effort and expense for other endangered species.

## Methods

### Sampling and Data Generation

Larval tissue samples were collected over three consecutive breeding seasons in the late spring of 2013, 2014, and 2015 using seines and dipnets. We timed our sampling to occur when larvae were large enough to sample non-destructively with small tail clips (Polich, Searcy, & Shaffer, 2013). Tail tips were placed in 95% ethanol within 30 seconds of clipping, larvae were immediately released at the site of capture, and tail tips were stored at −80C until use. A handheld GPS unit was used to locate ponds in the field, and final spatial coordinates and areas of ponds were taken from tracings of Google Earth images from February 2007. We sampled larvae from multiple locations within each pond to randomly sample the genetic variation present.

DNA was extracted using a salt extraction protocol (Sambrook & Russell, 2001), diluted to 100 ng/μL, and sheared for 28 cycles (30s on, 90s off) using the “high” setting on a Bioruptor NGS (Diagenode). After shearing, samples were dual-end size selected to approximately 300–500bp using 0.8X-1.0X SPRI beads (Rohland & Reich, 2012). Libraries were prepared with between 419 and 2000 ng of starting input DNA using Kapa LTP library prep kit half reactions (Kapa Biosystems, Wilmington MA), dual-indexed using the iTru system (Glenn et al., 2016), combined into pools of 8 (500ng/library, 4,000ng total input DNA) and enriched using a MYcroarray (Ann Arbor, MI) biotinylated RNA probe set designed from 5,237 exons from unique genes from the California tiger salamander genome (McCartney-Melstad et al., 2016). Given the close phylogenetic relationships of all members of the tiger salamander complex (O’Neill et al., 2013; Shaffer & McKnight, 1996), we predicted that most of the probes would also capture the eastern tiger salamander homologs. A total of 30,000 ng of c0t-1 prepared from *Ambystoma californiense* was used for each capture reaction to block repetitive DNA from hybridizing with probes or captured fragments. Probes were hybridized for 30 hours at 60C, bound to streptavidin-coated beads, and washed four times with wash buffer 2.2 (MYcroarray). Enriched libraries were then amplified on-bead with 14 cycles of PCR, cleaned using 1.0X SPRI beads, and sequenced on three 150bp PE lanes on an Illumina HiSeq 4000.

### Reference Assembly

We built a reference assembly for read mapping and SNP calling using the Assembly by Reduced Complexity (ARC) pipeline (Hunter et al., 2015). Reads from the 10 samples that received the greatest number of reads were pooled and mapped to the 5,237 *A. californiense* targets using bowtie2 v.2.2.6 (Langmead & Salzberg, 2012). Pools of reads mapping to each of these targets were independently assembled using SPAdes v.3.11.0 (Bankevich et al., 2012), and the contigs assembled for each target then replaced their respective targets and another round of mapping was performed to these contigs. This process was repeated for 10 iterations to extend assembled targets several hundred bp in both directions from their central probe-tiled regions. Reciprocal best blast hits (RBBHs) were then found to represent each target locus using blast+ 2.2.31 (Camacho et al., 2009). The set of RBBHs was then blasted against itself to find similar regions among targets, which may be indicative of chimeric assemblies. The longest contiguous region within each RBBH that did not exhibit similarity to other RBBHs was chosen as the target in the final assembly.

### SNP Calling and Genotyping

Reads for all samples were trimmed to 150bp and adapters were trimmed using skewer 0.2.2 (Jiang, Lei, Ding, & Zhu, 2014). These trimmed reads were then mapped to the reference assembly using BWA-mem 0.7.15 (Li, 2013). Read group information was added to the aligned reads and PCR duplicates were marked using picard tools v2.13.2 (https://broadinstitute.github.io/picard/).

SNP calling and genotyping was performed according to GATK best practices (DePristo et al., 2011; Van der Auwera et al., 2013). First, a set of high-quality reference SNPs was generated to assess and recalibrate base quality scores within each sample. To do this, HaplotypeCaller from GATK nightly-2017-10-17 (McKenna et al., 2010) was run separately on each sample in GVCF mode followed by joint genotyping with GenotypeGVCFs. Then, any SNP that met any of the following criteria were filtered from the reference set: QD < 2.0, MQ < 40.0, FS > 60.0, MQRankSum < −12.5, ReadPosRankSum < −8.0, QUAL < 100. Similarly, any indel that failed any of the following criteria were also removed from the reference set: QD < 2.0, SOR > 10.0, FS > 60.0, ReadPosRankSum < −8.0, QUAL < 100. Base quality score recalibration (BQSR) was then performed at the lane level (three different platform units among all of the read groups) using GATK.

HaplotypeCaller in GATK was then used with on-the-fly BQSR to generate sample-level GVCF files that were jointly genotyped using GATK’s GenotypeGVCFs function. The same hard filters were then applied to the resulting VCF files, except that all SNPs with QUAL values above 30 (instead of 100) were retained. Genotype calls with phred-scaled quality scores under 20 were set to “missing” data, and SNPs with greater than 50% missing data were removed. Samples with missing data rates greater than 30% were also removed.

Given the extremely large genomes of ambystomatid salamanders (roughly 30GB) (Keinath et al., 2015; Licht & Lowcock, 1991), we were concerned about the possibility of including duplicated paralogous loci in our analyses. We attempted to correct for this by filtering out loci that contained excessive heterozygosity, as differences between true paralogs interpreted as homologs typically appear as highly heterozygous sites. We used VCFtools v.0.1.13 to calculate p-values for heterozygote excess for every SNP (Danecek et al., 2011; Wigginton, Cutler, & Abecasis, 2005). Target regions that contained at least one SNP with an excess heterozygote p-value below 0.001 were removed from the analysis. A set of SNPs was then generated by randomly choosing a single SNP from each target region that did not contain any excessively heterozygous SNPs, and we refer to this as the “linkage-pruned” dataset.

### Population Genetic Analysis

The presence of isolation by distance (IBD)—the relationship between geographic and genetic distance—was tested at both the individual and pond (population) levels. Individual genetic dissimilarity was calculated as one minus the percentage of all biallelic SNPs that were identical-by-state using the snpgdsIBS function in SNPRelate v1.6.4 (Zheng et al., 2012). The correlation and the significance of the relationship between these genetic distances and geographic distance was calculated using a Mantel test with 999,999 permutations in the R package vegan 2.4-0 (Mantel, 1967; Oksanen et al., 2016). Fst values were calculated using pairwise differences between sequences in Arlequin v3.5.2.2 (Excoffier & Lischer, 2010) with 100,172 permutations of the data to compute p-values. Adjustment for multiple testing was performed using the Benjamini-Yekutieli method implemented in base R (Benjamini & Yekutieli, 2001). A Mantel test was run with vegan 2.4-0 to determine the relationship between Fst/(1-Fst) and distance between ponds.

To characterize the level of genetic diversity present in tiger salamanders on Long Island, we calculated nucleotide diversity (π) across all sequenced individuals and for each pond after pooling samples across years. First, GenotypeGVCFs from GATK was run with the includeNonVariantSites flag, removing indels and discarding targets that exhibited clear heterozygote excess compared to Hardy Weinberg equilibrium expectations. Sites where fewer than 50% of individuals were genotyped at a threshold quality of 20 were discarded, and per-site π values (generated using vcftools v1.15) were summed and divided by the total number of qualifying genomic sites.

The linkage-pruned dataset was visualized using principal components analysis (PCA) in the R package SNPRelate v1.6.4 (Zheng et al., 2012). The first eight principal components were plotted with letters corresponding to collection sites (ponds). The proportion of the variance explained by each principal component was also extracted using SNPRelate v1.6.4.

To estimate the number of distinct population clusters in the data, ADMIXTURE v1.3.0 was run using the linkage-pruned dataset for K=1 to K=30 with ten different random number seeds (Alexander, Novembre, & Lange, 2009). Each replicate was subjected to 100-fold cross validation (CV), and CV errors were used to choose a “reasonable” set of K values. If the standard deviation of CV values for any K value overlapped the standard deviation of the best-scoring K value, we included it as a reasonable value for K.

Effective population sizes (Ne) for each pond were estimated using the linkage disequilibrium (LD) method in NeEstimator v2.01 with a minor allele frequency cutoff of 0.05 (Do et al., 2014; Hill, 1981). We calculated separate Ne estimates for all ponds pooled across years, and also for our pooled set of all 282 samples. LD-based estimates of effective population size from single cohorts (years sampled) represent the harmonic mean between the effective number of breeders (Nb) and the true effective population size (Ne) (R. K. Waples, Larson, & Waples, 2016). Alternatively, as the number of pooled cohorts approaches the generation length (the average age of parents for a cohort), LD-based estimators should approach the true Ne (R. S. Waples, Antao, & Luikart, 2014; R. S. Waples & Do, 2010).

Effective population size estimates using the LD method can be downwardly biased for multiple reasons. First, estimates may be biased when many loci are used due to physical linkage among loci, given that the method assumes the loci are unlinked (R. K. Waples et al., 2016). This effect is predictable, however, and can be corrected if the number of chromosomes or total linkage map length is known. An estimate of the linkage map length is available for the closely related axolotl, *Ambystoma mexicanum*, and we used this number (4200cm) to correct estimates of effective population size for dense locus sampling by dividing them by 0.9170819 (or −0.910 + 0.219 x ln(4200)) (Voss et al., 2011; R. K. Waples et al., 2016). LD-based estimates of effective population size can also be downwardly biased when analyzing mixed cohorts in iteroparous species such as *A. tigrinum*, although this bias decreases as the number of sampled cohorts approaches the generation length of the species (R. S. Waples et al., 2014; R. S. Waples & Do, 2010). Therefore, single-cohort estimates of Ne were further corrected by dividing dense-locus adjusted estimates by 0.8781801, the product of two equations from Table 3 of Waples et al. (2014) that use the ratio of adult lifespan (estimated at 7 years for the closely related *A. californiense*) to age at maturity (4 years, also in *A. californiense)* (Trenham, Shaffer, Koenig, & Stromberg, 2000) to compensate for the downward bias introduced by iteroparity: (1.103−0.245 * log(7/4)) * (0.485+0.758 * log(7/4)). We used linear regression to quantify the relationship between pond area and effective population size, using multi-year estimates of Ne when available.

### Impact of Roads

We were interested in assessing the degree that roads restrict movement on this landscape. First, traffic data were downloaded from the New York State Department of Transportation (https://www.dot.ny.gov/tdv). Straight lines were drawn between ponds, and the annual average daily traffic (AADT) values of road segments intersected by these lines were summed, yielding a pairwise distance matrix of traffic between ponds (upper diagonal, Table 2). These values were divided by 11,702, the AADT of New York State Route 25 in the region, to scale the environmental distance matrix into units of “New York State Route 25 equivalents crossed”. The contributions of geographic and environmental (road) distances to the observed genetic variation were calculated using the genetic covariance decay model implemented in the software BEDASSLE (Bradburd, Ralph, & Coop, 2013), which avoids the statistical problems inherent in partial Mantel tests (Guillot & Rousset, 2013). This method fits a model of exponential decay in genetic covariance as a function of geographic and environmental distance. The final result is an expression of the contributions of geographic and genetic distance to observed spatial genetic patterns in comparable units. The MCMC_BB function in BEDASSLE was run for 5 million generations on the linkage-pruned dataset and sampled every 5,000 steps.

## Results

### Sampling

We genotyped 283 *Ambystoma tigrinum* larvae from 17 ponds spread over an approximately 35 km^2^ area (Figure 1, Table 1). More than 1.9 billion 150-bp sequencing reads were generated from three Illumina HiSeq 4000 lanes across these samples (mean=6.8 million reads/sample, min=1.8 million reads, max=10.9 million reads).

**Figure 1:**
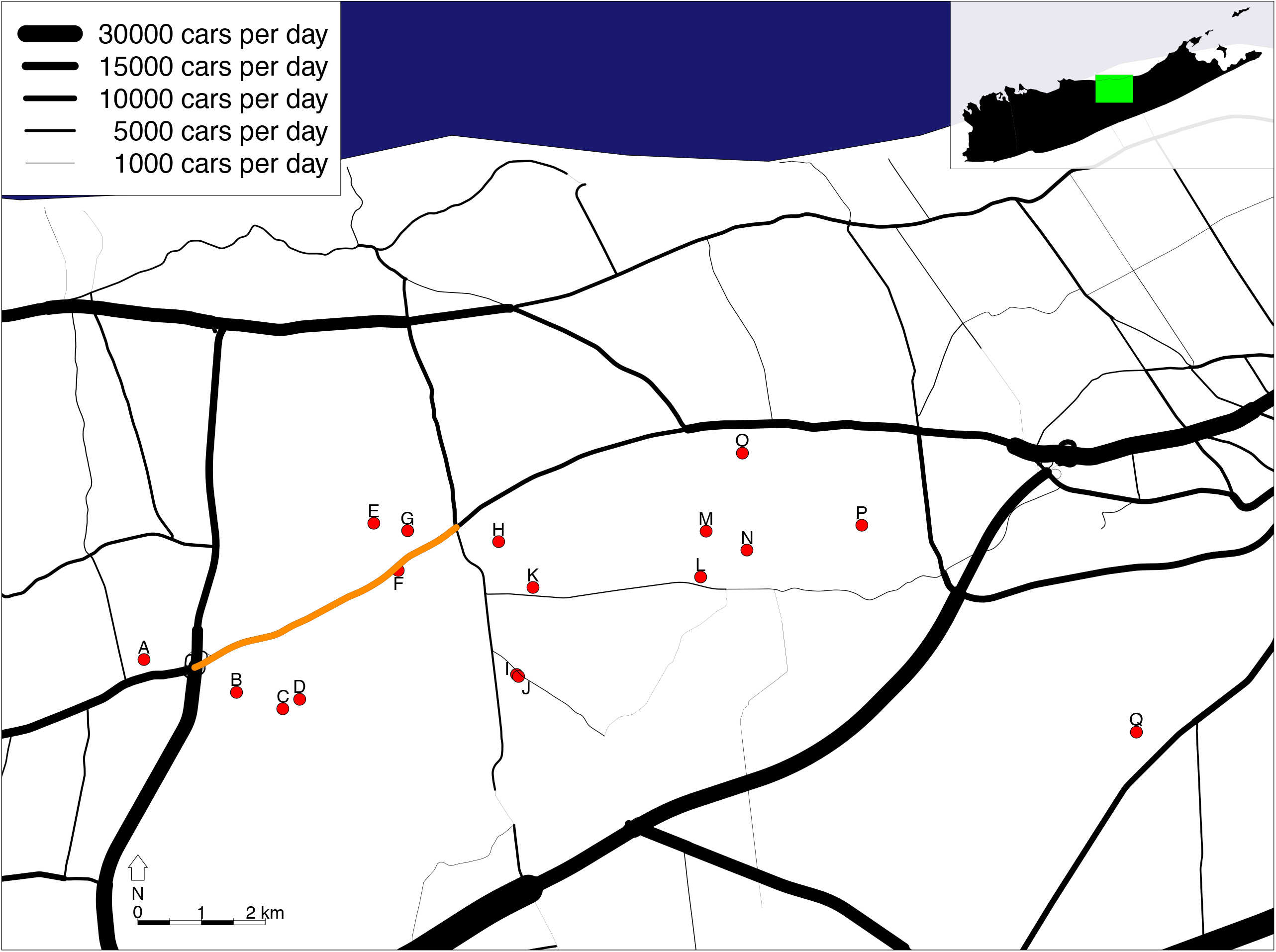
Map of sampling localities. Black lines indicate roads, and line thickness represents the average annual daily traffic (AADT) volumes of road segments. Sampling localities are represented by red dots. The road segment plotted in orange is the segment of New York State Route 25 used to standardize traffic measurements for BEDASSLE analyses (AADT=11,702).

**Table 1:** Pond localities, areas, π estimates, sampling, and effective population size estimates. Pond areas were estimated from Google Earth satellite images taken in February 2007. Single-year estimates were corrected for iteroparity-induced downward bias as explained in Methods, and both single-year and pooled-year estimates were corrected for dense locus sampling on chromosomes. Infinite values indicate that sample sizes were likely too small to estimate Ne—3 of 10, 10 of 10, and 10 of 10 replicates yielded infinite values for ponds I, L, and Q, respectively. N=number of samples included in analyses. Ne=Effective population size estimates using LD method, with standard deviations from 10 replicates. The “All” category denotes all 282 samples pooled across ponds.

| Pond | Latitude | Longitude | Pond Area (m<sup>2</sup>) | Nucleotide diversity ($\pi$) | N (2013/2014/2015) | $N_e$ +/- stdev[$N_e$] (years of sampling) |
| --- | --- | --- | --- | --- | --- | --- |
| **A** | 40.896379 | -72.892071 | 2147 | $1.79 \times 10^{-3}$ | 37 (10/18/9) | 11.47 +/- 0.11 (3) |
| **B** | 40.891766 | -72.874854 | 6413 | $2.03 \times 10^{-3}$ | 20 (0/10/10) | 39.54 +/- 0.69 (2) |
| **C** | 40.889497 | -72.866932 | 1706 | $2.05 \times 10^{-3}$ | 10 (0/0/10) | 14.03 +/- 0.26 (1) |
| **D** | 40.891043 | -72.863908 | 2039 | $1.93 \times 10^{-3}$ | 9 (0/0/9) | 19.25 +/- 0.75 (1) |
| **E** | 40.915705 | -72.849554 | 2493 | $2.01 \times 10^{-3}$ | 28 (10/9/9) | 27.57 +/- 0.36 (3) |
| **F** | 40.908597 | -72.845109 | 2898 | $2.06 \times 10^{-3}$ | 20 (10/0/10) | 31.62 +/- 0.56 (2) |
| **G** | 40.914317 | -72.842938 | 2094 | $1.97 \times 10^{-3}$ | 10 (0/0/10) | 17.98 +/- 0.43 (1) |
| **H** | 40.912580 | -72.826168 | 1840 | $2.13 \times 10^{-3}$ | 28 (10/10/8) | 92.36 +/- 1.38 (3) |
| **I** | 40.893704 | -72.823658 | 944 | $2.14 \times 10^{-3}$ | 5 (3/2/0) | Inf (2) |
| **J** | 40.893182 | -72.823465 | 3418 | $2.11 \times 10^{-3}$ | 30 (10/10/10) | 27.28 +/- 0.58 (3) |
| **K** | 40.906296 | -72.820671 | 10773 | $2.09 \times 10^{-3}$ | 29 (10/8/11) | 138.20 +/- 3.37 (3) |
| **L** | 40.907237 | -72.787736 | 8587 | $2.50 \times 10^{-3}$ | 1 (1/0/0) | Inf (1) |
| **M** | 40.913165 | -72.787206 | 7020 | $2.05 \times 10^{-3}$ | 8 (0/8/0) | 75.31 +/- 4.77 (1) |
| **N** | 40.910430 | -72.779946 | 464 | $1.97 \times 10^{-3}$ | 10 (10/0/0) | 9.91 +/- 0.25 (1) |
| **O** | 40.924112 | -72.780170 | 4710 | $2.06 \times 10^{-3}$ | 15 (10/5/0) | 63.92 +/- 2.44 (2) |
| **P** | 40.913681 | -72.758595 | 4854 | $2.01 \times 10^{-3}$ | 17 (10/0/7) | 38.59 +/- 0.87 (2) |
| **Q** | 40.883585 | -72.708374 | 1302 | $1.82 \times 10^{-3}$ | 5 (0/0/5) | Inf (1) |
| **All** | NA | NA | NA | $2.11 \times 10^{-3}$ | 282 (94/80/108) | 85.07 +/- 1.49 (3) |

### Reference assembly

The ten samples that received the most sequencing reads were pooled to generate a *de novo* reference assembly, for a total of 67 million merged and paired-end sequencing reads (11.7 billion total bp). Assembly of target regions with the ARC assembler produced a set of 61,621 contigs (39.9 million bp) from which 5,072 reciprocal best blast hits were recovered (6.6 million bp). After blasting these contigs against themselves, trimming self-complementary regions to the ends of contigs, and re-determining reciprocal best blast hits, a 6.5 million bp assembly with 5,068 target regions (96.8% of the originally targeted regions) was recovered for mapping reads and calling SNPs.

### SNP Calling and Genotyping

An average of 29.05% of raw reads mapped to the reference assembly using BWA-mem across all 283 samples (sd=2.50%, min=20.19%, max=34.13%). Removing PCR duplicates (read pairs that map to the exact same position on the reference, indicating that they may be PCR amplicons from the same molecule) resulted in an average of 16.95% unique reads mapped to the reference (sd=2.42%, min=8.57%, max=22.57%). After joint genotyping, a total of 79,233 raw SNPs were recovered across 4,414 target regions. Applying hard filters to SNP loci, setting the minimum genotype call quality to 20, discarding variants genotyped in fewer than half of all samples, and removing the one sample with a missing data rate greater than 30% yielded a total of 22,513 retained SNPs across 3,640 target regions. Tests for Hardy Weinberg equilibrium revealed 540 targets contained at least one SNP with clear (p<0.001) heterozygote excess, which is consistent with (though not definite evidence of) the presence of an unknown paralogous copy of this gene in the genome. We removed these target regions from the analysis, leaving a total of 12,955 biallelic SNPs across 3,095 target regions. The final matrix contained 282 individuals with a mean missing data rate of 8.09% (max=28.65%, min=2.13%, sd=4.53%), and the linkage-pruned dataset contained one random biallelic SNP from each final target for a total of 3,095 variants.

### Genetic variation and isolation by distance (IBD)

Nucleotide diversity within ponds ranged from 1.79×10^−3^ to 2.50×10^−3^ (Table 1) and was 2.11×10^−3^ for the combined 282 samples. IBD was apparent at both the individual and pond levels. Mantel tests of the significance of correlations between individual and pond genetic and geographic distances yielded p-values of 1×10^−6^ and 1.2−10^−5^ and Mantel R^2^ statistics of 0.1743 and 0.4092, respectively. This strongly indicates that there is a significant relationship between geographic and genetic distance, even at the extremely fine scale studied here.

Pairwise Fst values between ponds ranged from 0.005 to 0.210 (136 comparisons, median=0.064, sd=0.042, Table 2). Using Benjimini-Yekutieli (BY) corrected p-values, 118 out of 136 of these pairwise comparisons were significant. Of the 18 non-significant pairwise comparisons, 16 were from pond L, which contained only a single sample and therefore had extremely low power. Many of the highest Fst values were from pairwise comparisons containing ponds A or Q. These ponds are both geographical outliers separated by greater geographic distance (pond Q) and by more major roads (both A and Q) from other sites than are all other ponds (Figure 1).

**Table 2:** Pairwise Fst values between ponds (lower diagonal) and cumulative annual average daily traffic (AADT) distances between ponds (upper diagonal—in units of New York State Route 25 traffic equivalents). Bolded Fst values are not significant (p > Benjamini-Yekutieli-corrected 0.05).

| Pond | A | B | C | D | E | F | G | H | I | J | K | L | M | N | O | P | Q |
| --- | --- | --- | --- | --- | --- | --- | --- | --- | --- | --- | --- | --- | --- | --- | --- | --- | --- |
| A | * | 3.798 | 4.035 | 3.838 | 1.127 | 2.789 | 1.740 | 3.118 | 3.683 | 3.683 | 3.283 | 3.283 | 3.118 | 3.283 | 3.118 | 3.283 | 6.191 |
| B | 0.126 | * | 0.000 | 0.000 | 1.000 | 0.000 | 1.000 | 0.329 | 0.310 | 0.310 | 0.494 | 0.494 | 0.494 | 0.494 | 0.329 | 0.494 | 2.818 |
| C | 0.131 | 0.019 | * | 0.000 | 1.000 | 0.000 | 1.000 | 0.329 | 0.310 | 0.310 | 0.494 | 0.494 | 0.494 | 0.494 | 0.494 | 0.494 | 2.818 |
| D | 0.128 | 0.030 | **0.022** | * | 1.000 | 0.000 | 1.000 | 0.329 | 0.310 | 0.310 | 0.494 | 0.494 | 0.494 | 0.494 | 0.494 | 0.494 | 2.818 |
| E | 0.147 | 0.067 | 0.069 | 0.071 | * | 1.000 | 0.000 | 1.329 | 1.425 | 1.425 | 1.329 | 1.329 | 1.284 | 1.329 | 1.284 | 1.284 | 3.905 |
| F | 0.119 | 0.043 | 0.045 | 0.049 | 0.040 | * | 1.000 | 0.329 | 0.310 | 0.310 | 0.329 | 0.329 | 0.329 | 0.329 | 0.329 | 0.329 | 2.905 |
| G | 0.173 | 0.086 | 0.087 | 0.091 | 0.050 | 0.060 | * | 1.329 | 1.425 | 1.425 | 1.329 | 1.329 | 1.329 | 1.329 | 1.284 | 1.329 | 3.905 |
| H | 0.116 | 0.038 | 0.040 | 0.042 | 0.051 | 0.022 | 0.064 | * | 0.262 | 0.262 | 0.000 | 0.000 | 0.000 | 0.000 | 0.000 | 0.000 | 2.577 |
| I | 0.139 | 0.051 | 0.062 | 0.061 | 0.068 | 0.035 | 0.087 | 0.036 | * | 0.000 | 0.262 | 0.262 | 0.262 | 0.262 | 0.262 | 0.262 | 2.508 |
| J | 0.121 | 0.048 | 0.051 | 0.046 | 0.055 | 0.030 | 0.078 | 0.026 | 0.023 | * | 0.262 | 0.262 | 0.262 | 0.262 | 0.262 | 0.262 | 2.508 |
| K | 0.112 | 0.036 | 0.039 | 0.040 | 0.046 | 0.018 | 0.060 | 0.005 | 0.033 | 0.020 | * | 0.000 | 0.000 | 0.000 | 0.000 | 0.000 | 2.577 |
| L | **0.163** | **0.073** | **0.064** | **0.101** | **0.072** | **0.040** | **0.112** | **0.020** | **0.046** | **0.042** | **0.026** | * | 0.000 | 0.000 | 0.000 | 0.000 | 2.481 |
| M | 0.152 | 0.067 | 0.070 | 0.080 | 0.073 | 0.047 | 0.095 | 0.038 | 0.057 | 0.052 | 0.039 | **0.031** | * | 0.000 | 0.000 | 0.000 | 2.485 |
| N | 0.175 | 0.090 | 0.101 | 0.105 | 0.102 | 0.083 | 0.125 | 0.066 | 0.088 | 0.075 | 0.065 | **0.070** | 0.067 | * | 0.000 | 0.000 | 2.485 |
| O | 0.128 | 0.044 | 0.049 | 0.056 | 0.055 | 0.030 | 0.076 | 0.022 | 0.038 | 0.033 | 0.019 | **0.034** | 0.032 | 0.061 | * | 0.000 | 3.382 |
| P | 0.135 | 0.062 | 0.063 | 0.070 | 0.066 | 0.042 | 0.091 | 0.039 | 0.052 | 0.044 | 0.037 | **0.033** | 0.054 | 0.077 | 0.031 | * | 3.382 |
| Q | 0.210 | 0.129 | 0.142 | 0.146 | 0.138 | 0.118 | 0.164 | 0.112 | **0.136** | 0.115 | 0.108 | **0.159** | 0.133 | 0.151 | 0.112 | 0.117 | * |

### Principal Component Analysis

The first eight principal components (PCs) together account for 13.5% of the total genetic variation (Figure 2). PC1 groups samples from pond A to the exclusion of the other samples, while PC 2 does the same for samples from ponds E and G. PC 3 separates ponds B, C, and D from the other ponds (especially ponds N and part of J), and PC4 appears to be an axis of variation between ponds J and Q/N. Finally, PCs 5, 6, 7, and 8 correspond to axes that differentiate ponds N, P, and Q, along with some samples from ponds A and J. Overall, clustering of single ponds and small groups of closely adjacent ponds is quite apparent, indicating the presence of easily detectable population structure with these genomic data.

**Figure 2:**
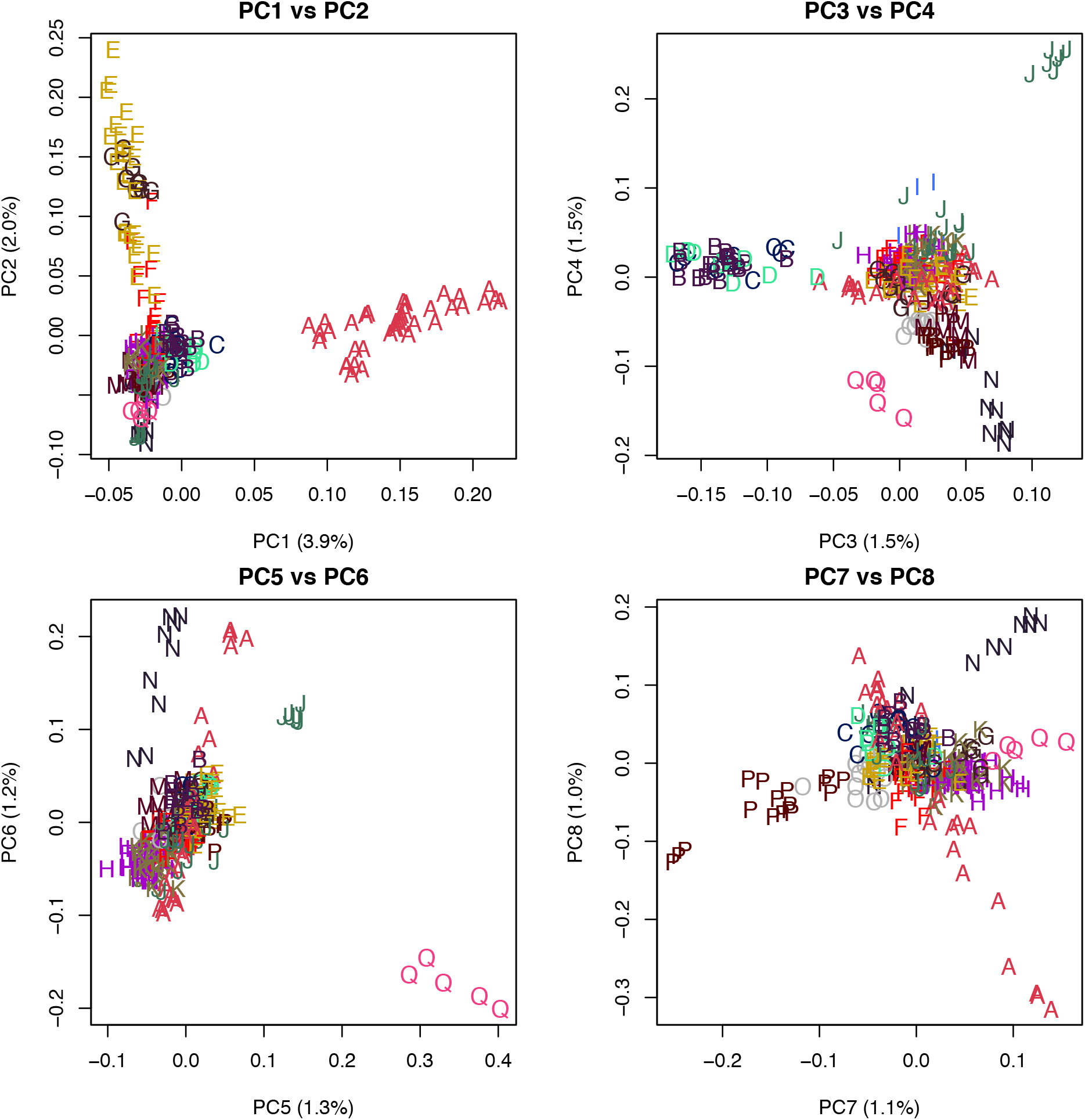
First eight principal components of the data. Letters on the graph correspond to samples from the same pond. Colors are used only to aid in distinguishing between letters.

### Population Clustering

The value of K in ADMIXTURE with the lowest mean CV error was K=12. Three other K values (9, 10, and 11) had CV error standard deviations that overlapped with K=12 (Figure S1). Admixture proportions for K=9 through K=12 are shown in Figure 3, and are split by both pond and sampling year (Glasbey, van der Heijden, Toh, & Gray, 2007). Results from ADMIXTURE analyses corroborated the qualitative patterns observed in the PCA. Pond A generally formed one to three clusters to the exclusion of all other ponds (as recapitulated in PCs 1, 6, and 8). Ponds B, C, and D formed a single cluster to the exclusion of other ponds (as also seen in PC 3). Similarly, ponds E and G form a unique cluster at K=9 (corresponding to PC 2), but are separated into their own private clusters at K=10 through K=12. Pond F, geographically separated from its closest neighbors (ponds E and G) by NY State Route 25, appears strongly admixed at K=9 through K=12. Ponds H, I, J, K, L, and M appear to be strongly associated across all K values (although ponds I, L, and M appear highly admixed), except for one year of sampling in pond J (2014), which produced a group of animals that formed their own cluster. Pond N is distinct across all K values (which can also be seen on PCs 3–8). Pond O appears highly admixed across all K values, but tends to share a considerable admixture component with the cluster formed by pond P (and pond Q for K=9 and K=10). At K=11 and K=12, pond Q forms its own cluster to the exclusion of all other ponds, a pattern that is also apparent in PCs 5 and 6.

**Figure 3:**
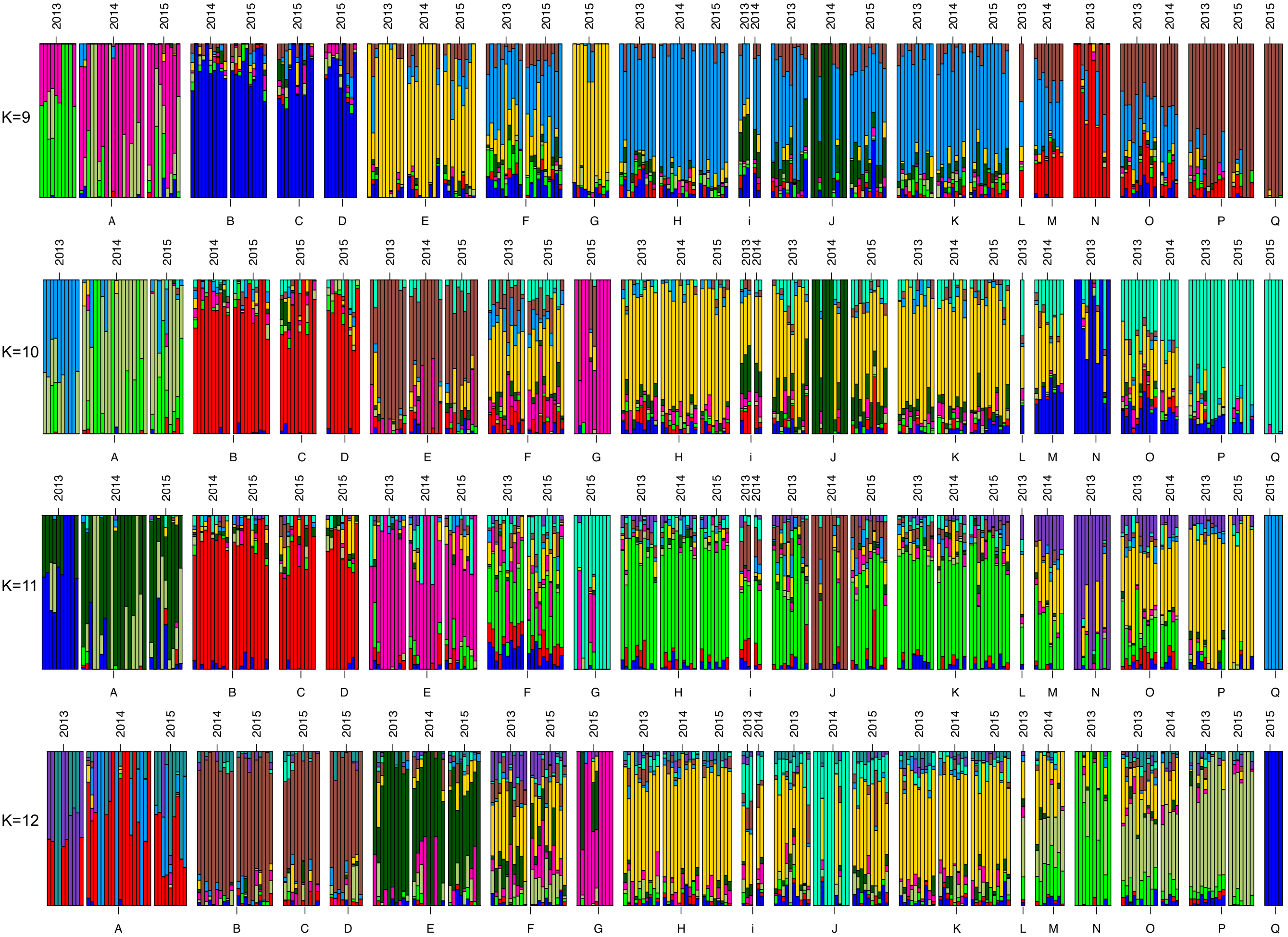
Admixture results from all 282 samples. Letters correspond to ponds from the sample map (Figure 1). Thin white spaces separate sampling years within ponds, and thicker white spaces separate ponds from one another.

### Effective Population Size

Estimates of effective population size ranged from 9.81 for pond N to 138.20 for pond K (Table 1). Pond surface areas were strongly correlated with effective population size estimates (p=0.00122, R^2^=0.5623, Figure 4). Dropping pond H, the clear outlier in Figure 4, increases the significance of this relationship (R^2^ of 0.849, p<0.000005), while dropping both ponds H and K (the extreme outlier in terms of pond size, and therefore a highly influential value) still returns a significant positive relationship (R^2^ of 0.671, p<0.0007). The number of samples included in the calculation of Ne was not correlated with the resulting Ne estimate (linear regression p=0.2448, adj R^2^=0.03341), suggesting that sample size was not a driver of Ne estimates.

**Figure 4:**
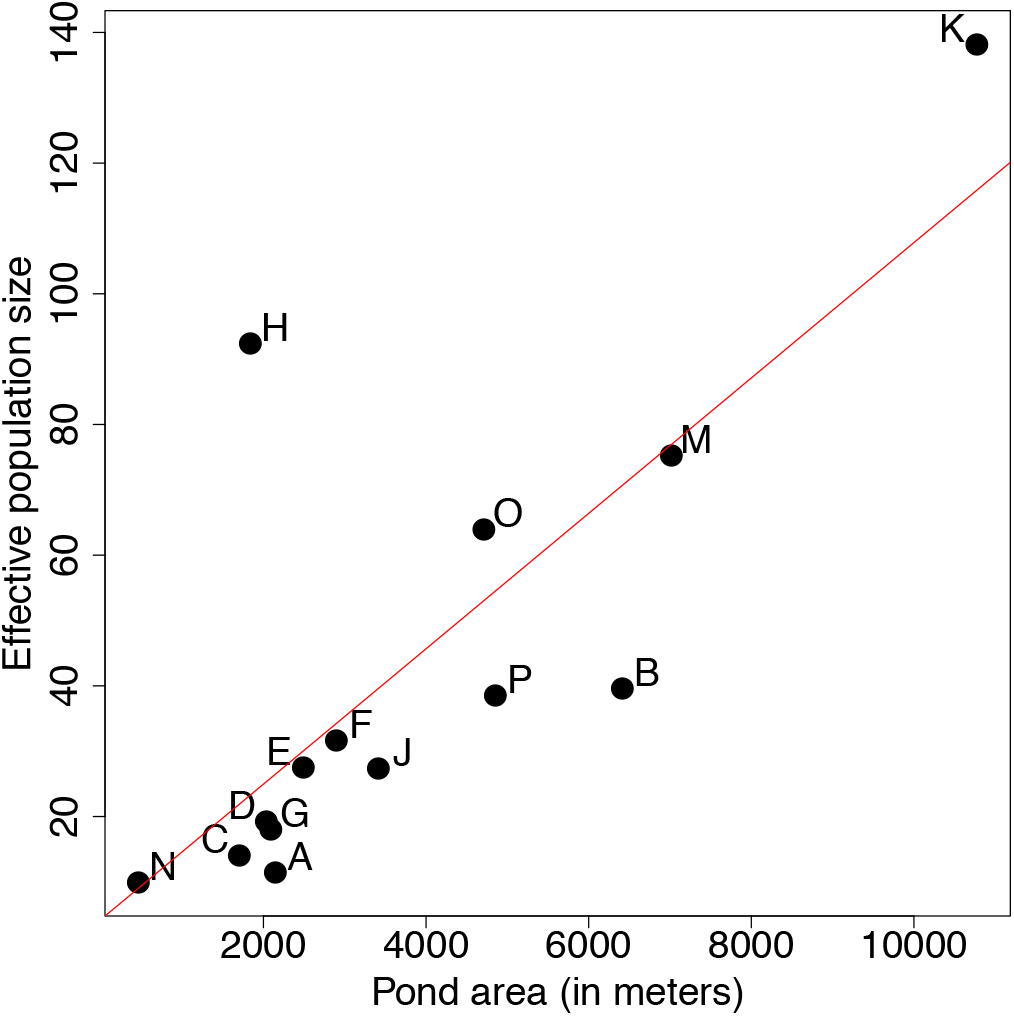
Relationship between pond area and effective population size estimate (p=0.00122, adjusted R^2^=0.5623). Ne estimates represent multiple-cohort calculations if multiple cohorts were sampled, otherwise single-year estimates are used. Ponds I, L, and Q were omitted because they did not contain enough samples to generate non-infinite Ne estimates across replicates.

### Roads as Barriers to Dispersal

Roads play a strong role in structuring among-pond genetic divergence in Long Island tiger salamanders. We calculated the relative contributions of geographic distance and roads (scaled by traffic) between ponds with 10 independent runs of the covariance decay model in BEDASSLE. Nine out of 10 runs found that the presence of one road with the same traffic as New York State Highway 25 (orange line in Figure 1) has an equivalent effect on population divergence as 6.2km to 8.8km of Euclidean distance, while the remaining run measured this effect at 4.4km (posterior medians in Figure 5). Posterior predictive sampling (Figure S2) indicated that this model did a good job of modelling the effects of geographic distance and isolation by roads on pairwise genetic relationships among ponds. These results indicate that dispersal is severely limited by roads, and that human activity has contributed to isolation of ponds in this region.

**Figure 5:**
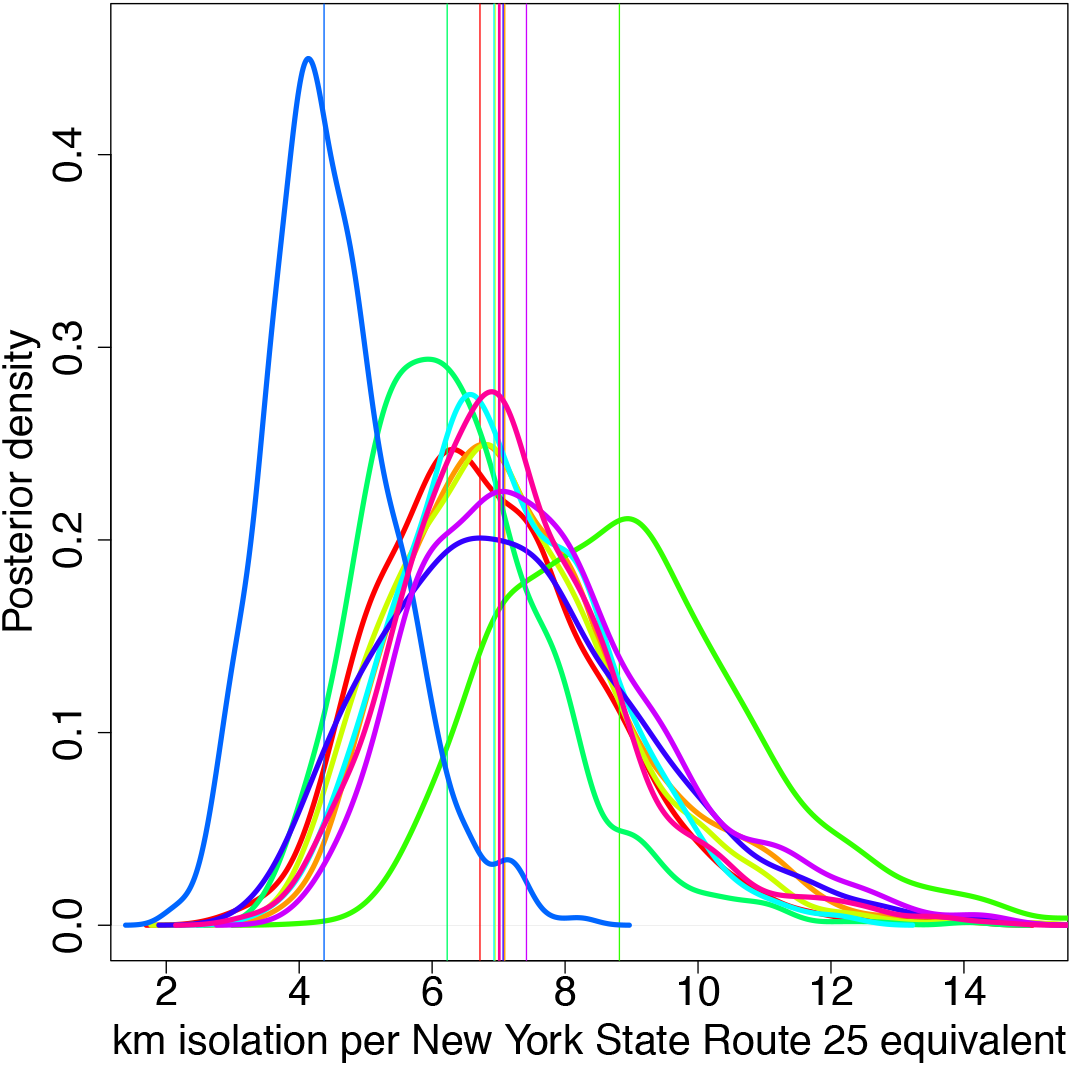
Posterior distributions of BEDASSLE aE/aD values with 100 generations of burnin. Each curve represents 1 of 10 independent BEDASSLE runs, and vertical lines indicate median values of the posterior distributions. The x-axis quantifies the relative impacts of geographic and environmental distance. For instance, a value of 7 indicates that a road with traffic equal to New York State Route 25 isolates ponds by the equivalent of 7km of geographical distance.

## Discussion

Population genetic structure is often difficult to detect and quantify accurately among subtly differentiated populations, including those in close geographic proximity (Wright, 1943). As conservation geneticists, however, we are often interested in understanding limitations to gene flow imposed by recent anthropogenic activities. A satisfactory explanation of the factors that impinge on gene flow requires a clear picture of how populations are structured on the landscape. Our RADseq approach provided the resolution necessary to evaluate the influence of human disturbance on the movement of endangered salamanders, and in particular the role that roads play in limiting among-pond dispersal. Posterior distributions of geographical and environmental coefficients calculated in BEDASSLE showed that the presence of two moderately sized roads isolated ponds by the same amount as roughly 14 km, or nearly the entire length of the study system. In contrast to microsatellites, genomic-scale data provided the resolution required to detect the effects of geographic distance and roads on dispersal, despite the relatively recent construction of roads in the area. Assuming that this effect on dispersal accumulates with time, it also suggests that roads are probably more important than geographic distance in shaping population structure, given the magnitude of their effect over only a few generations. In support of this, a previous study of the yellow-spotted salamander (*Ambystoma maculatum*) in central New York state found between 50% and 82% adult mortality due to car strikes at a road crossing (Wyman, 1991), levels that are more than sufficient to cause the measurable impacts on gene flow that we document (Gibbs & Shriver, 2005). This knowledge will be critical for future conservation planning.

Prior studies have illustrated that amphibian genetic differentiation is sometimes (e.g. Savage et al., 2010; I. J. Wang, 2009, 2012, I. J. Wang et al., 2011, 2009), but not always (e.g. Jehle, Burke, & Arntzen, 2005; Lampert, Rand, Mueller, & Ryan, 2003; R. A. Newman & Squire, 2001; Zamudio & Wieczorek, 2007) detectable at very fine spatial scales. The key question, for molecular ecology and conservation biology alike, is whether such results represent biological differences among species and habitats or the resolving power of the molecular markers utilized, which itself is shaped by deeper-time demographic processes such as bottlenecks and range expansions (Slatkin, 1993; Watterson, 1984). While microsatellite loci have been extremely valuable for conservation genetics, a panel of 20 microsatellites has been shown to be approximately as effective for estimating genetic relationships as just 50 SNP loci (Santure et al., 2010). Increasing the number of microsatellite loci above 20 is both difficult and expensive, but it is now straightforward to scale the number of SNPs assayed into the (tens of) thousands, increasing our ability to distinguish barriers to gene flow that are subtle or have only been operating for relatively few generations (Anderson et al., 2010; Patterson et al., 2006). As genomic-scale datasets become comparable with microsatellites in terms of cost and feasibility, the added resolution from thousands of loci will give a particular boost to population genetic studies in systems with low genetic diversity, and will open entire new classes of analyses to both low- and high-diversity systems.

The current study is among the first to use thousands of nuclear loci across hundreds of individuals in a large-genome (∼ 30 gigabase) amphibian and represents an opportunity to directly compare results between two genetic approaches in the same system. While little genetic clustering was apparent in the microsatellite loci analyzed by Titus et al. (2014), our dataset of thousands of nuclear SNPs revealed clear population genetic structuring among the same breeding ponds of *Ambystoma tigrinum* on Long Island. The major genetic patterns in our data are readily apparent in both ADMIXTURE and PCA results, and our genetic analyses of pond structuring returned generally consistent results across years (Figure 3), suggesting that the observed patterns of genetic structure do not represent idiosyncratic interannual variation.

Species with low genetic diversity require collecting data from a greater number of genetic loci to detect population structure (Patterson et al., 2006), and one cause of low genetic diversity is a recent range expansion. Church *et al.* (2003) analyzed *Ambystoma tigrinum* mitochondrial DNA and determined that New York was likely recolonized by salamanders from Pleistocene refugia in North Carolina. This was corroborated by Titus et al. (2014), who found low genetic diversity in microsatellite loci in New Jersey and Long Island tiger salamanders. To better understand whether this low genetic diversity was responsible for the low resolving power of microsatellite (Titus et al. 2014) compared to target capture (current study) datasets, we calculated genetic diversity for each pond and across all 282 samples. Nucleotide diversity across the 282-sample dataset was 2.11×10^−3^, indicating that Long Island tiger salamanders have lower diversity than 67 of the 76 species measured by Romiguier *et al.* (2014). Such low levels of genetic diversity may well be a function of Pleistocene glacial history and recolonization dynamics, and generally indicate that large genetic datasets are necessary to discern real, but subtle population structure as occurs in this system.

Breeding ponds that we examined generally exhibited small effective population sizes (< 100), consistent with results found for many other amphibian species (McCartney-Melstad & Shaffer, 2015; Phillipsen, Funk, Hoffman, Monsen, & Blouin, 2011; Schmeller & Merilä, 2007). Our estimates (mean=43.4) are larger than, but of the same magnitude as microsatellite-based estimates recovered by Titus *et al.* (2014) using the sibship method (J. Wang, 2009), which had a mean value of 20.9. We did, however, recover several ponds with effective population sizes higher than 44, which was the maximum value recovered by Titus *et al.* (2014). These included pond H (Ne=92.4), pond K (Ne=138.2), pond M (Ne=75.31), and pond O (Ne=63.92), and may indicate that the area around these ponds, which was not directly sampled by Titus et al. (2014), harbors greater effective population sizes than elsewhere on Long Island.

We also recovered a clear relationship between pond size and effective population size (p=0.00122, R^2^=0.5623, Figure 4), a critical result for two reasons. First, this area/population size relationship has been previously reported in two independent studies of another member of the tiger salamander complex, *A. californiense* (I. J. Wang et al., 2011; I. J. Wang & Shaffer, 2017); the consistency across studies indicates that the largest breeding ponds harbor the greatest potential for species persistence genetically and demographically. In the absence of genetic data, managers working across diverse habitats (CA, NY) may be able to use pond size as a reasonable proxy for population size, and therefore, conservation value. Second, these data emphasize the importance of connectivity and metapopulation dynamics in this system. The pond for which surface area did the poorest job predicting Ne, Pond H, had a much higher effective population size estimate than predicted (Figure 4). Pond H is geographically closest to Pond K, which has the largest effective population size estimate in the study. The landscape between Pond H and Pond K is forested with no major roads or other anthropogenic barriers to gene flow, the Fst value between ponds H and K is the lowest of any pairwise comparison between ponds (Fst=0.005, Table 2), and these ponds were consistently recovered in the same ADMIXTURE cluster (Figure 3). Taken together, this provides strong evidence that migration is common between Ponds H and K, and that the effective population size of Pond H is augmented by its close relationship with the very large Pond K. Given the importance of large Ne for current and future adaptation and resiliency (Frankham et al., 2017), this confirms the critical importance of maintaining connectivity and metapopulation dynamics for these endangered populations.

## Conclusion

From a biological perspective, our results demonstrate that endangered *Ambystoma tigrinum* on Long Island generally have small to modest effective population sizes that are correlated with the surface area of ponds, naturally limited migration among ponds, and severely constrained dispersal and disrupted metapopulation dynamics because of roads. The interrelationships between these factors are important for conservation management. Small effective population sizes imply that ponds are more likely to suffer random demographic extinction, and highly structured populations indicate that locally extirpated ponds (such as those that do not fill with water for many years in a row) may not be easily recolonized by individuals from nearby sites even a kilometer or two distant. Roads, especially those with high traffic, and other human activities add to these natural dynamics, emphasizing the critical importance of conserving blocks of contiguous habitat with pond complexes that can persist as semi-isolated metapopulations. Within this Long Island landscape, there are several pond clusters that periodically share migrants (ponds B, C, and D; ponds H, I, J, K, L, and M; and ponds O and P). Maintaining these dynamics is likely critical for the long-term persistence of these locally endangered tiger salamanders because they are potential source populations for nearby ponds as they go locally extinct and require rescue. However, the presence of roads disrupts this pattern, as seen by the tendency of nearby ponds separated by major roads to fall into different genetic clusters (such as Pond A vs. ponds B, C, and D and Pond F vs. ponds E and G).

From a methodological perspective, our work demonstrates that a genomic approach was critical to detect and quantify existing, fine-scale population structure in this post-glacially recolonized area. Titus *et al.* (2014) recovered little genetic structure among endangered populations of Long Island tiger salamanders and inferred relatively high migration rates between ponds. Our genomic approach revealed the opposite pattern, with restrictions in movement between many groups of ponds despite low overall levels of genetic differentiation. The generality of such drastic differences in inferred biological processes is an unanswered empirical question that requires additional comparative studies across taxa, landscapes, and demographic histories.

Finally, from a conservation perspective, our genomic results suggest that ponds that are not separated by major roads almost certainly have increased resilience to local extirpation via demographic rescue from neighboring ponds, and that they also may benefit from genetic rescue as climate and other anthropogenic changes necessitate rapid evolutionary responses. Our strongest conclusion is that efforts should be made to prevent activities that restrict movement among clusters of ponds, particularly as decisions on conservation and property use are made in this critical open space. The landscape bounded by the Brookhaven National Laboratory to the west and Pond Q to the east represents a relatively intact, ∼100 km^2^ ecological oasis and the last stand for this endangered amphibian in the northeastern US. It should remain that way.

## Acknowledgments

Funding was provided by the Andrew Sabin Family Foundation. EMM and HBS also received support from NSF-DEB 1457832 and NSF-DEB 1257648. We thank Alvin Breisch, Andy Sabin, Pete Davis, and Jeremy Feinberg for assistance with fieldwork, Megan Sha and Rice Zhang for laboratory support, and Kirk Lohmueller and Beth Shapiro for helpful comments on an earlier version of this manuscript. This work used the Vincent J. Coates Genomics Sequencing Laboratory at UC Berkeley, supported by NIH S10 Instrumentation Grants S10RR029668 and S10RR027303. Computing resources were provided by XSEDE (Towns et al., 2014) and the Comet supercomputer at SDSC, supported by NSF ACI-1548562.

---

**Figure S1:**
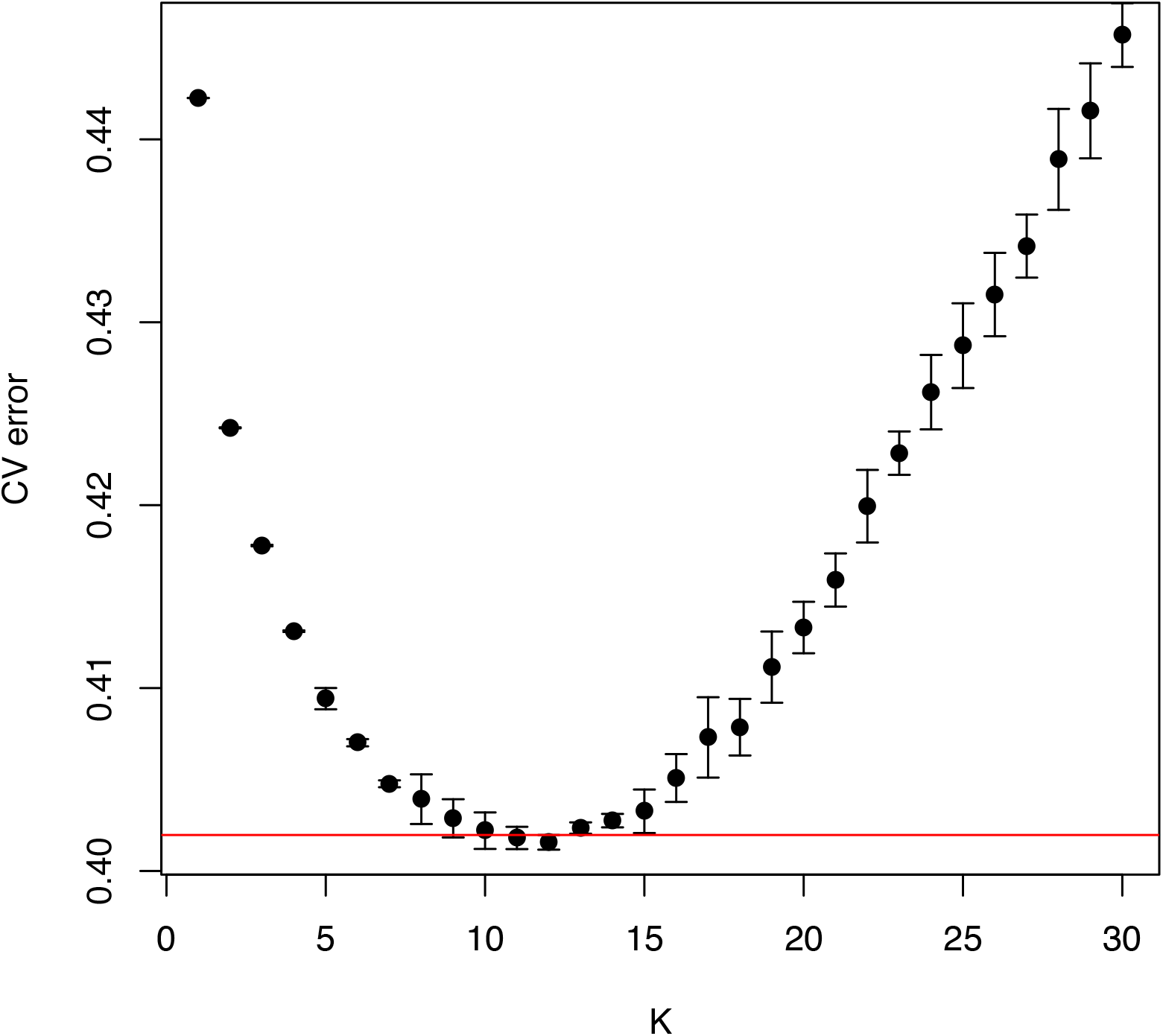
Cross-validation error means and standard deviations from 10 ADMIXTURE runs using different seeds. The red line is drawn at the mean+SD of the best-performing K value (K=12). The standard deviations for K=9 through K=12 overlap this line.

**Figure S2:**
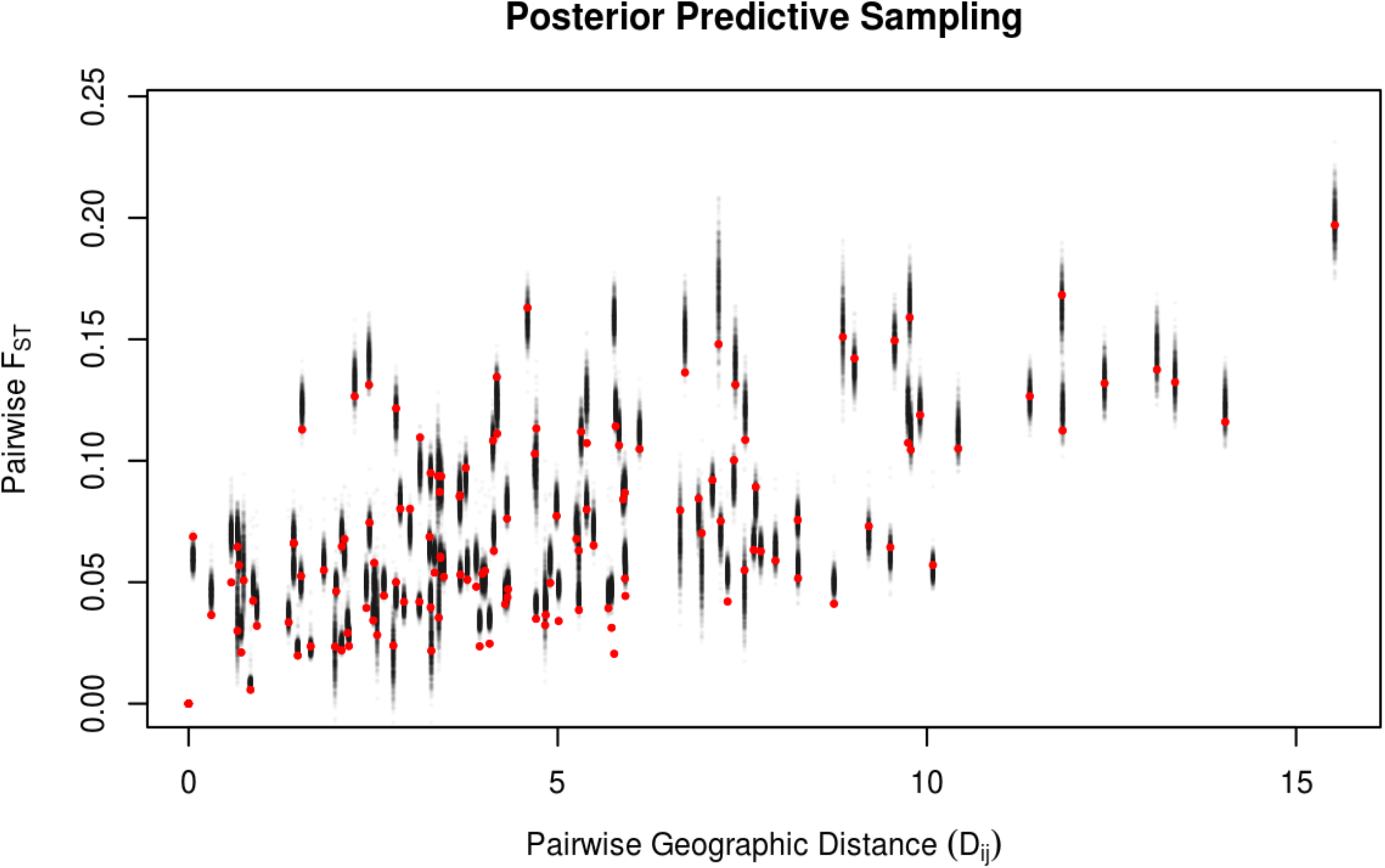
Visualization of model fit of highest-probability BEDASSLE run. Red dots indicate empirically observed Fst values between ponds, while black dots indicate predicted Fst values from 1000 posterior predictive samples.

